# Adaptive Phenotypic Plasticity for Life History and Less Fitness-Related Traits

**DOI:** 10.1101/367284

**Authors:** Cristina Acasuso-Rivero, Courtney J. Murren, Carl D. Schlichting, Ulrich K. Steiner

## Abstract

Organisms are faced with variable environments and one of the most common solutions to cope with such variability is phenotypic plasticity, a modification of the phenotype to the environment. These modifications influence ecological and evolutionary processes and are assumed to be adaptive. The assumption of adaptive plasticity allows to derive the prediction that the closer to fitness a trait is, the less plastic it would be. To test this hypothesis, we conducted a meta-analysis of 213 studies and measured the plasticity of each reported trait as coefficient of variation (CV). Traits were categorised according to their relationship to fitness into life-history traits (LHt) including reproduction and survival related-traits, and non-life-history traits (N-LHt) including traits related to development, metabolism and physiology, morphology and behaviour. Our results showed, unexpectedly, that although traits differed in their amounts of plasticity, trait plasticity did not correlate with its proximity to fitness. These findings were independent of taxonomic groups or environmental types assessed and raise questions about the ubiquity of adaptive plasticity. We caution about generalising the assumption that all plasticity is adaptive with respect to evolutionary and ecological population processes. More studies are needed that test the adaptive nature of plasticity, and additional theoretical explorations on adaptive and non-adaptive plasticity are encouraged.

## INTRODUCTION

Adaptation to varying environments has long been a central question in ecology and evolution [1]. In times of global change and increased habitat fragmentation due to anthropogenic activities, individuals and populations experience novel environments more frequently than in the recent past. Therefore, they are confronted with increased environmental variability and less predictable spatial and temporal environments [2]. Populations can deal with such varying conditions by either local adaptation or phenotypic plasticity [3,4]. Local adaptation involves genetic differentiation specific to each environment; whereas phenotypic plasticity allows single genotypes to express different phenotypes under diverse environmental conditions. Such tracking of environmental change via plasticity can be achieved with or without genetic differentiation [3,4]. The role that plasticity plays for ecological and evolutionary processes is controversial in the context of speciation and diversification, adaptation to novel environments, population viability, population management, and the invasion of species [5–7]. The controversy arises because phenotypic plasticity is a potential mechanism for responding to environmental challenges [5,9,10], but on the other hand, might also hinder adaptation to novel environmental conditions [11].

Many of these controversies rest upon the assumption that phenotypic plasticity is adaptive [10,12,13], though contrasting findings have been presented [14]. Studies reveal different outcomes depending on traits, species and environments. Previous approaches to assessing the frequency of adaptive plasticity have been based on analyses of reciprocal transplant studies [15], transcriptome and proteome analyses [see references in 14], or meta-analyses comparing plastic and canalized responses, particularly in plants [14]. To date, no standard approach across organisms has been accepted for how to assess whether plasticity in a specific trait is adaptive or not. Well-documented cases of adaptive plasticity have been reported, as well as conditions under which plasticity is disadvantageous or maladaptive [15–18]. However, because fitness is frequently defined through traits more closely related to fitness [life-history traits (LHt)], and different traits respond in similar ways to environmental changes, such traits can be correlated and thus confounded with environmental effects [14,19]. For instance, being larger is often correlated with higher fitness, but larger sizes are also commonly enabled by nutrient rich environments. Yet we often lack data on what the optimal size in such a rich environment would be and if it is realised [20,21]. Even powerful approaches such as reciprocal transplant experiments do not circumvent the challenge of determining optimal phenotypes.

Under the assumption that phenotypic plasticity is adaptive, LHt should show less plastic responses than N-LHt. In order to test this prediction, we used an indirect approach by reviewing 24 years of research publications, covering a wide range of environmental conditions and spanning taxonomic groups. Our approach provides power to the study of phenotypic plasticity by generalising the question and adding and comparing information from a great variety of sources. In the next section we outline the arguments of why this prediction can be used as a condition that plasticity is adaptive. We then present and discuss the results of our meta-analysis.

### Plasticity in Life-History and Non-Life-History Traits

Phenotypically plastic traits evolve as any other quantitative trait by natural selection acting on genetic variation among genotypes; here genotypes vary in their reaction norms, the level of phenotypic plastic responses across environments [22]. Selection for plasticity is strong under environmental conditions when organisms can rely on environmental cues; and tracking of phenotypes to the environment increases relative fitness. If, however, environmental cues are not reliable, selection may favour fixed genetically determined traits (environmental phenotypic canalization), and phenotypic plasticity may not be possible or beneficial [3]. To this end, selection acts to optimise the trait expression in each environment to maximise fitness in that environment or across environments [23].

Since trait plasticity does not evolve differently than other traits, plastic traits more closely related to fitness are predicted to be under stronger selection for genetic canalization, both within and across environments [24–28]. So, genetic variability in reaction norms will erode faster in LHt (here considered as reproduction or survivorship) compared to N-LHt (all other traits, that are less directly related to fitness) [24,27,28]. However, the difference in selection strength for genetic canalization does not predict whether LHt should exhibit more or less plasticity than N-LHt. In the next section, we illustrate how such a prediction can be derived from evolutionary theories of plasticity and demographic theory [29,30] related to similar arguments discussed in previous studies [31,32].

An adaptively plastic genotype, considered as a generalist, expresses a phenotype that more closely matches the fitness optimum across environments than a less plastic genotype, a specialist; the latter has high fitness in one environment but not in others [22,33]. This foundational theory of the evolution of plasticity leads to the fundamental expectation that fitness across environments should vary more for specialists that lack plasticity in key morphological, physiological, or behavioural traits compared to adaptively plastic generalists when phenotypic (fitness) optima differ among environments. Considering that fitness components respond in an integrative manner across traits and environments, for individuals sharing the same genotype, it is expected that LHt should vary less across environments compared to N-LHt. To put it another way, because of trait integration, trade-offs, and optimisation of fitness across traits and environments, N-LHt are expected to moderate and buffer – through their plasticity – the effect of environmental variation on LHt. We use this prediction to test whether plasticity might be adaptive or not.

Related arguments rooted in demographic theory have been formulated by Caswell [31]. He extends previous developed models of homoeostasis [34] to a demographic setting [35,36]. The population growth rate *λ* serves as fitness and he argued that plasticity in morphological, physiological, or behavioural traits may reduce variance in *λ*, whereas plasticity in life history traits should lead to increased variance in *λ*, as long as life history traits are not negatively covarying.

Extensions of these foundational demographic theories sustain aspects of these arguments. For instance, the demographic buffering hypothesis is rooted in stochastic demographic theories that state that vital rates (mortality and fertility) of population projection models with higher sensitivities (with respect to *λ*) should be negatively correlated to variability in these vital rates across environments. Empirical evidence for the demographic buffering hypothesis is mixed. In the context of our study, it is crucial to note that the demographic buffering hypothesis does not predict whether one trait type is expected to express higher plasticity. This lack of prediction is likely because the theoretical focus and empirical tests of the demographic hypothesis have been on LHt, and not on N-LHt, i.e., age-specific survival or reproduction [37]. The environmental canalization hypothesis (distinct from genetic canalization) states that the potential demographic impact of fitness components and their temporal variability, like plasticity, should be negatively correlated based on investigations from age-structured populations [38]. This hypothesis has been empirically tested by comparing variance in survival among juvenile versus adult individuals and it is supported for longer-lived mammals but not shorter-lived ones [38].

The underlying arguments of both the demographic buffering and the environmental canalization hypotheses are derived from stochastic population theory [39,40]. This theory illustrates how stochastic population growth rate, *λ*_s_, is reduced by high variability in vital rates among different environments experienced across time. Focal vital rates are fitness components such as reproduction and survival, which are the LHts we discuss above. Following this, an increase in fitness can be achieved simply by reducing variability in reproduction and survival across time or environments, without increasing either mean survival or reproduction. Adaptive phenotypic plasticity is predicted to do exactly this: high plasticity of non-life history traits (N-LHt) buffer the environmental effects on fitness, leading to reduced variance in traits closely related to fitness (also referred as LHt) [31,32].

## METHODS

In order to test whether LHt exhibit lower plasticity than N-LHt, we conducted a meta-analysis on studies reporting reaction norms in different trait types across environments and species. We performed this meta-analysis investigating 24 years of research literature on phenotypic plasticity. We selected papers employing the keywords “life history & morphology & plasticity” between 1991-2006 from all databases in *Web of Knowledge (WoK-ISI)* and between 2007-2011 from *Web of Science (ISI)*. We set out with 583 papers in total, including a few studies known to us that met the criteria published between 1987 and 2011. From this initial set, we extracted data from 213 studies reporting reaction norms. Quantification of the reaction norms was achieved through the computation of the CV of the trait expression across environments, a dimensionless parameter allowing the evaluation of proportional responses as a mean-standardised measure [41]. We computed 5,885 coefficients of variation including 211 species exposed to at least two and at most 11 environment levels reporting phenotypic plasticity. As a visual validation for reporting and publication bias [42], we present a plot of the number of studies through time and a funnel plot in the supplementary material (Fig. S1 and S2).

Traits were categorised as: Life-history traits (LHt): survival and reproduction; and non-life history traits (N-LHt): behaviour, morphology, metabolism and physiology, and development. When available we extracted information directly from tables or from the text, alternatively we extracted information from figures using the software *ImageJ* [43]. 1). Environmental variation was grouped into six categories: 1) Environment Quality, 2) Interspecific Interactions, 3) Intraspecific Interactions, 4) Intrinsic Resources, 5) Photoperiods and Light; and 6) Temperature. When genotypes or families were included, taxonomic groups were classified based on species ID as defined by the NCBI taxonomy database [44]. We clustered organisms at the taxonomic level of *Phylum* (Taxa in Table 1; Table S1 in the supplementary material describes additional information extracted and its categorisation where relevant).

We used Mixed-Effect Models (lme4) in software R [46; R Core team, version 3.2.3] for analyses, formulating models on log transformed CV as the response variable and using reference ID as a random effect to account for confounding effects within the same publication. We also used the number of data points retrieved from a single study — *Repetitions* (Table S1) — as weights to assess potential bias towards a few studies or species. The focus of our study was the evaluation of the explanatory variables *Trait* type (including the 5 broad categories), *Environment type*, and *Taxonomic* group. Other factors listed in Table S1 were explored but only marginally contributed to the observed variance (analyses not shown). We used information theoretical approaches (Aikake Information Criterion: AIC) [46] to select among our candidate models and interpreted a model to be better supported for any ΔAIC≥2.

To gain a better quantitative understanding of how frequently our prediction was met, we computed, for each pair of LHt and N-LHt, the difference in CV within studies (n=3,939), limiting our analysis to the 103 studies that measured both categories of traits. Negative values indicate that the N-LH trait showed higher plasticity, supporting our hypothesis, and positive values reveal the opposite.

Finally, we tested the assumption that life history traits are genetically more canalized compared to non-life history traits [24–26]. Unfortunately, reaction norm plots generally do not indicate genotypes with unique labelling, preventing distinguishing replicates from truly different genotypes (strains, families, populations). For this analysis on genetic canalization, we excluded studies that compared groups (“genotypes”) that are likely genetically not very distinct (e.g., workers and drones in bees, experimental evolution studies, studies that compared same genotypes or populations over time (season, years), or where sex was the only difference recorded). We included distinct populations, different strains, and subspecies that shared the same NCBI identity but were noted by the authors of the studies as dissimilar. Using this subset of data, we estimated within each study, for each genotype, and each trait the mean CV, i.e. mean plasticity for each genotype/strain, assuring that each genotype is equally weighted. From the mean CVs, we estimated the variance among genotypes in their reaction norms. To test if LHt were more genetically more canalized than N-LHt, we used a generalised linear model with a Gamma error structure (lme4) to distinguish variance in CV between LHt and N-LHt,

## RESULTS

The 5,885 CVs quantifying the plastic responses to environmental variation across 211 eukaryote species, comprising 59% Invertebrates, 34% Chordates, 6% Plants and 1% Green Algae. Details on data and taxonomic identities are shown in the supplementary material 1 (DRYAD: doi:10.5061/dryad.72s8g4j).

Substantial variation in plasticity (CV) was revealed within groups and among groups of trait types, taxonomic groups and environment type (Table 1, Fig. 1, Fig. S3). The plastic response to environmental variation (CV) when only a single factor was assessed, was best explained by trait type rather than environment type or taxonomic group. A null model (intercept only model), did not perform better than any of the single factor variable models. The differences in plasticity among trait types are not correlated to how closely traits are associated with fitness (Fig. 1). Adding taxa as an additional variable to the model with trait type explained more variability than adding environment type (Table 1). Interactive effects between trait types and environment type or taxonomic group did further improve the model fit (Table 1), but make biological interpretation challenging. In contrast to our hypothesis, none of these models showed a clear correlation between plasticity (CV) and how closely traits are connected to fitness (Fig. 1). Hence, we show that the phenotypic response to environmental variation is dependent on the interaction between trait and environment type, but with no relation to how close the trait is to fitness.

**Table 1:** Competing linear models selected based on AIC. Models used as response variable the log transformed Coefficient of Variation (scaled plasticity), explored combinations of trait type, environment and taxa as fixed effects, and included reference study as a random effect as well as repetitions (sample size) as weighting factor.

| | Model | AIC | ΔAIC |
| --- | --- | --- | --- |
| **Null Model** | Intercept only | 19494 | 546.41 |
| **1** | Trait type | 19111.41 | 163.82 |
| **2** | Environment | 19463.77 | 516.18 |
| **3** | Taxa | 19476.06 | 528.47 |
| **4** | Trait type + Environment | 19076.17 | 128.58 |
| **5** | Trait type + Taxa | 19083.74 | 136.15 |
| **6** | Trait type * Environment | 18970.24 | 22.65 |
| **7** | Trait type * Taxa | 19044.05 | 96.46 |
| **8** | Trait type + Environment + Taxa | 19051.65 | 104.06 |
| **9** | **Trait type \* Environment + Taxa** | **18947.59** | **0** |

**Figure 1:**
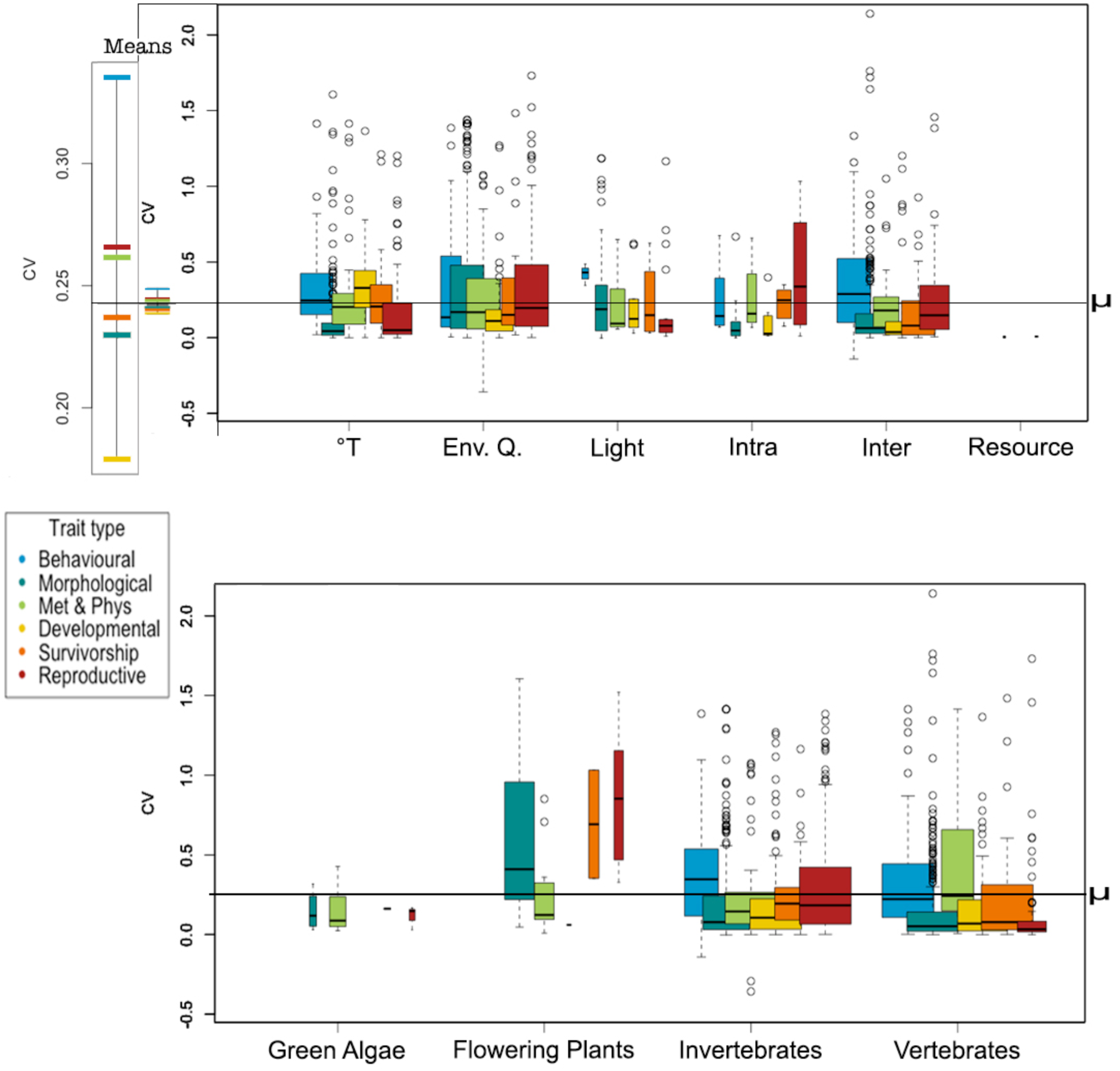
Plasticity (logCV) among different trait type, environment type and taxa. Trait types are ordered roughly in accordance with relative distance to fitness. We categorised life-history traits (LHt) into survivorship and reproductive; and non-life-history traits (N-LHt) into behavioural, morphological, metabolism and physiology, and developmental. The top left side panel shows the trait means across all environments on an expanded scale. To its right are these means shown at the same scale as of the boxplot. Top right panel, coefficients of variation (CVs) for each environment and trait type (colour coded). The widths of the boxes are relative to the amount of data; whiskers represent the standard deviation. The grand mean (μ) is denoted by the black line across all graphs. °T: Temperature. Env. Q.: Environmental Quality. Light: Photoperiods and light. Intra: Intraspecific interactions. Inter: Interspecific interactions. Resource: Intrinsic resources. Lower panel: CV among taxa and trait types. The widths of the boxes are relative to sample size; whiskers represent the standard deviation. For green algae and plants, no behavioural or developmental data are available.

Plants expressed high plasticity (CV) and they showed limited evidence of increased CV of LHt compared to N-LHt, contradicting our main hypothesis (Fig. 1). Also, Plants was the taxonomic group that drove most of the difference among the taxa. Analyses that excluded the plant data, found no difference between taxonomic groups, an interaction between trait type and environment, yet with no relationship to how closely a trait is related to fitness (Supplementary material 5).

Our results were robust to explorations of other factors - the type of experiment (lab or field), whether natural populations or more controlled genotypes were used, how many environments per study were explored, the number of data points per study, the study year, or whether only life-history traits, only non-life history traits or both types of traits were reported within a stud (Table S2). None of these changed the results qualitatively or had much quantitative influence. A lack of influence of these other factors suggests that our sample size is sufficiently large, and that our results are robust and general across different types of study systems.

When we compare pairs of life-history and non-life history traits within studies, i.e. estimating the difference in CV between a LHt and a N-LH trait, we see that our hypothesis was supported as often as it was rejected (Fig. 2). Negative values, supporting our expectation of adaptive plasticity (i.e. non-life history traits exhibiting higher plasticity), were equally common as positive values (interpreted as revealing non-adaptive plasticity). For most comparisons, differences in CV were close to zero (Fig. 2), indicating that life history and non-life history traits within studies show similar plasticity, even though we see general differences in plasticity among trait types (Fig. 1). Behavioural traits might tend to express more non-adaptive plasticity (positive differences like metabolism and behaviour with reproduction), while morphological traits might tend to express more adaptive plasticity (negative differences like metabolism and survival).

**Figure 2:**
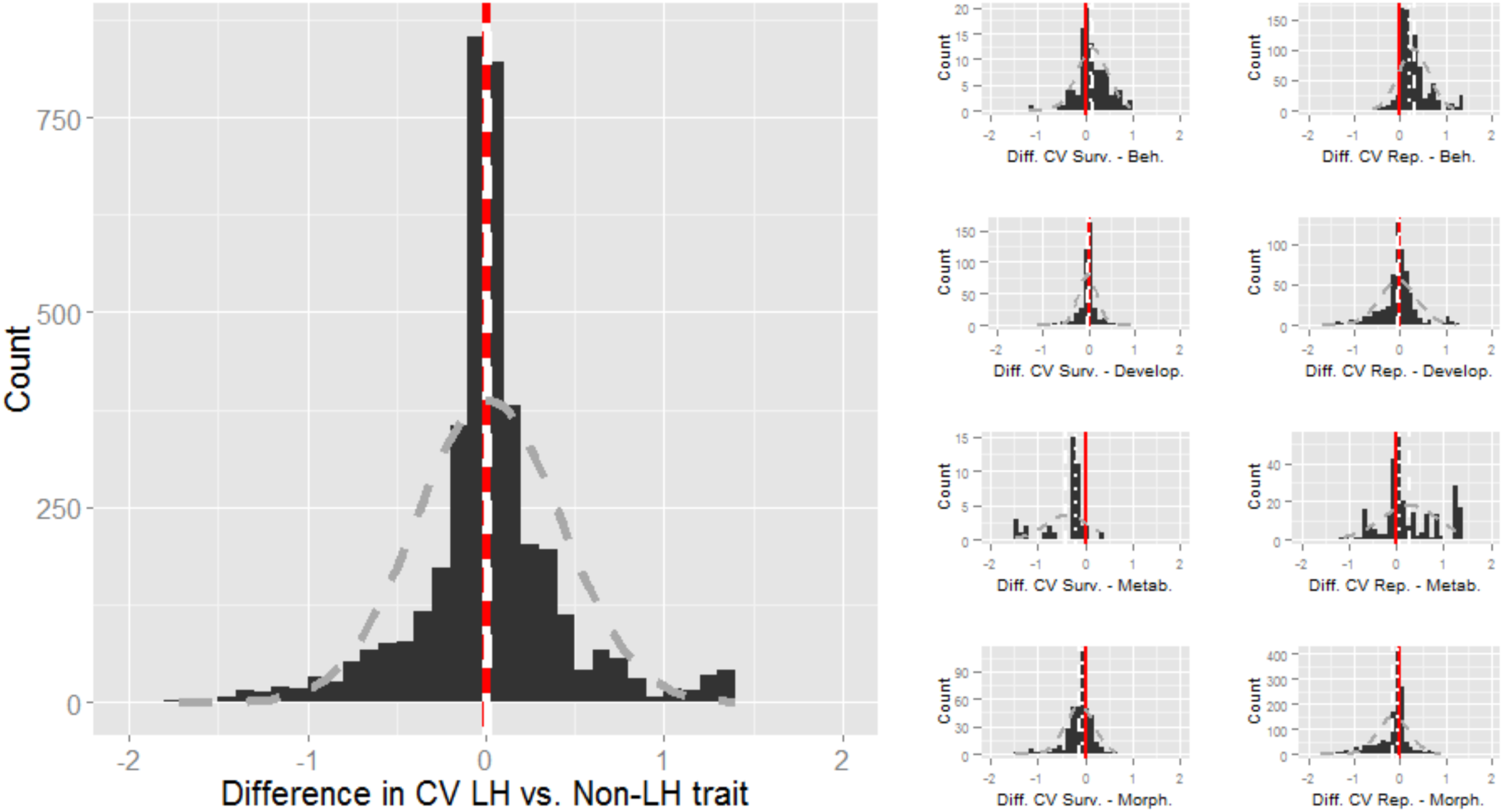
Distribution of differences in CV between life history traits and non-life history traits (CV_LHt_^_^CV_N-LHt_), large panel. Small panels illustrate distributions of differences in CV between survivorship and non-life history traits (Behavioural, Developmental, Metabolism and Physiology, Morphological) left column of panels, and reproductive and non-life history traits (right column of panels). Red solid line depicts no difference in CV (0), white dashed line depicts the mean, dotted line depicts the median. The grey dashed line depicts a normal distribution based on the observed mean and variance.

Even though LHt did not show lower plasticity (Fig. 1), we still expected that LHt should be genetically more canalized than N-LHt. In contrast to our expectation, LHt survival and reproduction, did not show lower levels of variability in reaction norms among genotypes, compared to N-LHt (Fig. 3). We found significant differences in the variance among genotypes in their reaction norms, with behavioural traits showing the highest variance among genotypes, but developmental, physiological, and morphological traits showed less variance in their reaction norms compared to the life history traits. Thus, there is no evidence for increased genetic canalization for LHt compared to N-LHt.

Note that the variability in genetic canalization (Fig. 3) was not linked to the mean level of plasticity exhibited (Fig. 1) but remember Fig. 3 is only based on a subset of data allowing no direct comparison.

**Figure 3:**
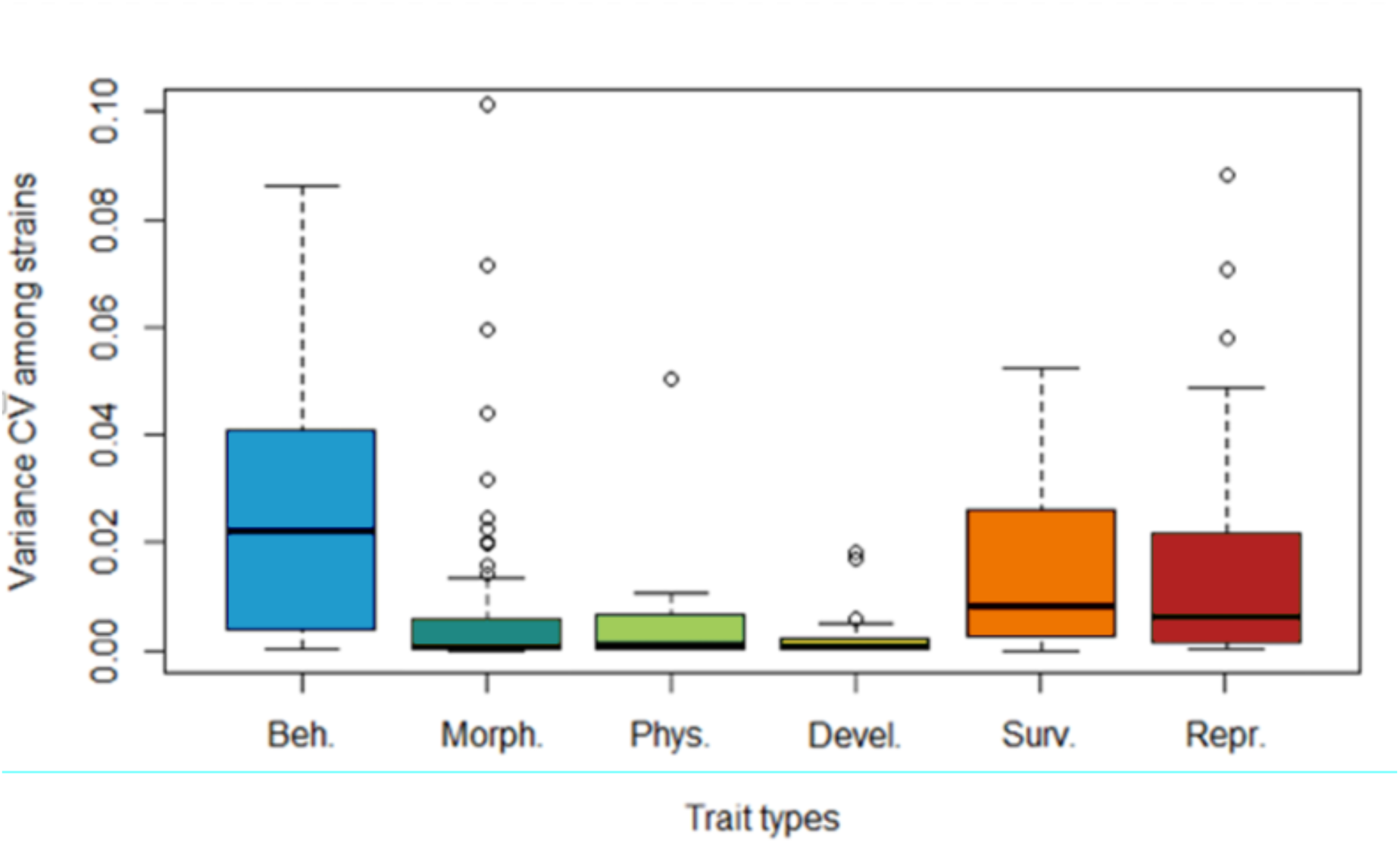
Variance in CV among genotypes within the same study and trait, plotted for the different trait types: Beh: Behaviour, Morph: Morphology, Phys: Metabolism and Physiology, Devel: Development, Surv: Survival, Repr: Reproduction. Note y-axis is limited to values <0.1 for better visibility, i.e. not all outliers are shown. Model selection for generalised linear models with a Gamma error structure and Variance of CV as response variable: AIC for model with trait types (-1426.6), null model (intercept only model) AIC (-1422.0).

We report a negative correlation between the overall CV of each trait type (Fig. 1), as a measure of plasticity, and the CV among genotypes (Fig. 3), as an inverted measure of genetic canalization (r=0.77, Spearman; Fig. 4). This suggests that less plastic traits will also be the ones that are fixed faster into the populations. However, as illustrated by figure 4, we show that the relationship between canalization and plasticity is not related to the distance from fitness. We found, as expected, that behavioural traits were more plastic and the least canalized. But we also found that morphological and developmental-related traits are the least plastic and more canalized than the other trait types. Our results also illustrate that reproductive and metabolic-related traits showed the least canalization and exhibited only a mid-plastic response when compared to other trait types. Traits related to survival showed an intermediate plastic response as well as an intermediate canalization.

**Figure 4:**
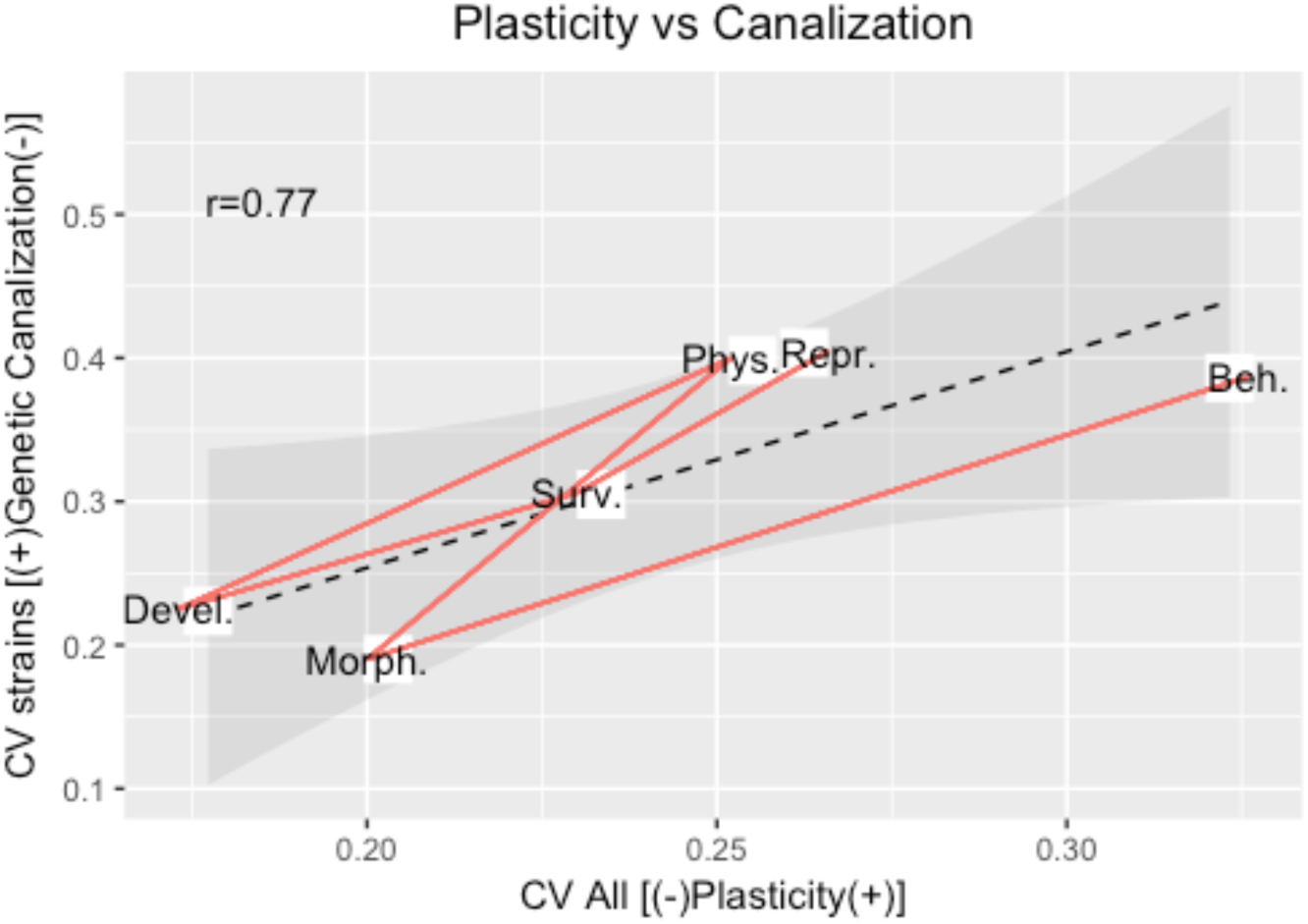
The correlation between plasticity and genetic variance, as an inverted measure for genetic canalization, is significant (r=0.77, Spearman). But the order of the traits did not correspond to their proximity to fitness. The black dashed line illustrates the linear relation between the trait types if ordered as follows: Developmental, Morphological, Survivorship, Metabolism & Physiology, Reproduction, and Behaviour. The red line links the traits in order as defined previously in relation to their distance to fitness: Beh-Morph-Phys-Devel-Surv-Rep. If the distance to fitness was a main determining factor to the relationship between plasticity and genetic canalization, both lines should coincide.

## DISCUSSION

Our results show that trait types did not correlate, as predicted, with how closely the traits were related to fitness but did differ in their amount of plasticity. Taken together, life history traits (LHt) are not less plastic than non-life history traits (N-LHt); and LHt are not buffered against environmental variation by N-LHt. Our results suggest that non-adaptive or potentially maladaptive responses in plasticity might be common [47]. The meta-analytical techniques we used, allowed us to aggregate information producing robust findings independent of the larger taxonomic groups, study conditions and types, and excluding the influence of potential publication biases (Fig. S2). Hence, our results were robust and seem general. Our results are in line with previous studies raising the question of the ubiquity of adaptive plasticity and highlight the challenges in differentiating between adaptive and non-adaptive plasticity [14]. The patterns we reveal suggest that identifying adaptive processes from phenotypic plasticity studies may be beyond the reach of studies that focus only on one or few traits and one or a few species or populations. We stress a need for caution on assumptions that plasticity is adaptive and suggest a re-evaluation of conceptual work that may have assumed this. We also encourage further explorations about the role non-adaptive plasticity.

To critically reflect on our results that plasticity is not necessarily adaptive, we revisit the arguments behind our hypothesis. Our central argument is based on basic evolutionary theories that suggest that the strength of selection varies with trait type, such that different selective forces should lead to different evolutionary outcomes [24–26]. Our first assumption is that survivorship and reproductive traits are more closely related to fitness than other traits. Fitness, the population growth rate *λ*, is made up of fitness components, and both, the traits that we define as LHt as well as the other traits, can be seen as such components. Although N-LHt probably also influence fitness indirectly through their influence on reproduction and survival, the influence of LHt is more direct and therefore stronger [48]. For example, body size influences both survival and reproduction in many systems [49,50], but is not perfectly correlated with survival or reproduction [51], and is thus less closely related to fitness.

Following from the proximity to fitness of LHt, selection for genetic canalization is expected to be stronger, and thus should reduce variability in plasticity among genotypes [24-26]; we did not find such patterns (Fig. 3). It could be that higher genetic canalization might be observed within studies, but a comparison of levels of canalization of LHt and N-LHt within studies finds no more frequent lower variances in CV of LHt (results not shown).

The strong relation between trait plasticity and canalization suggests the more plastic a trait, the less canalized it will be, supporting previous theories. However, we found that the relationship to fitness is not necessarily what determines the level of canalization. The lack of association indicates that the difference in selective forces for genetic canalization does not determine whether traits closely related to fitness (LHt) exhibit more or less plasticity compared to N-LHt. According to our categorisation, the traits were more fixed and less plastic resulting in the following order: Developmental, Morphological, Survivorship, Metabolism & Physiology, Reproduction, and Behaviour. Interestingly, survival traits fell out in the middle of both the plastic and genetic canalization responses, which might argue for some kind of buffering role among traits.

We are by no means suggesting there are no trade-offs between traits but questioning them in respect to their relationship to fitness. Evolutionary theories of life histories rely heavily on trade-offs among LHt [52]. If LHt negatively covary with each other they could also buffer each other against environmental variation and thereby weaken the expected environmental buffering of N-LHt on LHt [31] (Caswell 1983). Supporting previous studies, we do not find negative correlations among LHt traits (Fig. S4).

Most studies in an evolutionary ecological framework aim at conditions that are comparable to natural conditions. Maladaptive plastic responses are predicted to occur under rare or novel environments when cryptic genetic variation is released [17,18]. While our results suggest that non-adaptive or potentially maladaptive plastic responses might be common (Fig. 2), we cannot evaluate to what extent the different studies employed novel environments. From our own empirical work and from other meta-analyses on plasticity [53], the inclusion of novel environments in plasticity studies is frequent but by no means universal. The more the studies that included novel environments, the more likely a bias against our hypothesis. A more general challenge that we face, similar to other studies on plasticity, is not knowing what the optimal plasticity is in a given set of environments. For example, a plastic response that overshoots an intermediate optimum will also be seen as maladaptive, and such overshooting in an N-LHt would support our hypothesis that N-LHt are more plastic. This becomes even more challenging to quantify empirically as natural environmental variation is changing rapidly through climate and land use change.

We realise there is, as for most studies, a potential bias caused by the selected methodology. For instance, the scaled plasticity (CV) could introduce noise, but it is an accepted method to standardise variability among traits with fundamentally different units [54]. Our sample size should be sufficiently large for generalising our overall findings, supported by the absence of a major publication bias (Fig. S1), although relevance of findings for specific groups might suffer from low amount of data (e.g. green algae). The results are robust to a number of other factors and do not depend on specific types of studies. These arguments give us confidence in our results, despite the highly variable and diverse patterns of plasticity detected. Certainly, we encourage others to follow up on these general understandings and patterns gained from our meta-analysis and continue to follow up with empirical studies of species for which we know that adaptive, non-adaptive (neutral) and maladaptive plasticity have been found.

Plants show the highest level of plasticity which could be related to them being largely sessile, lacking some types of opportunity to avoid environmental variation through habitat selection. Such limitation might lead to particularly strong selective forces for plasticity [55]. However, if this high level of plasticity would be adaptive, we would expect that plants exhibit less plasticity in life history traits compared to non-life history traits, but the reverse was observed (Fig. 1- Lower panel). A meta-analysis of results of plant reciprocal transplant experiments also found no differences in the relative plasticity of life history traits vs. morphological traits [14]. Their results suggest however that, for those traits that are plastic, putatively adaptive plasticity is more common than non-adaptive responses. Whether this pattern holds for other sessile organisms (e.g. certain fungi or marine invertebrates) is worthy of future targeted investigations.

Our results support growing evidence [14,56–58] that much plasticity might be neutral, or even maladaptive. Our study does not reveal the causes that prevent the evolution of adaptive plasticity, but there are many that can be posited: variability in environmental conditions is high enough that environmental cues might not be as reliable as assumed [32]; environmental frequencies do not select for plasticity as argued by others [59]; costs and limits of phenotypes inhibit plasticity evolution [60]. We are by no means suggesting that plasticity is necessarily non-adaptive, and our results do not suggest so, but we call for caution about the generalising assumption about adaptive plasticity.

## ACKNOWLEDGMENTS

We would like to thank the Bettencourt Foundation for its support. Special thanks to Alice Demarez, Josh Van Buskirk, Josh Auld, Anurag Agrawal, and Rick Relyea. Miguel Tejedo, Federico Marangoni, Cino Pertoldi, Alex Ritcher-Boix, Germán Orizaola, Alfredo G. Nicieza, David Álvarez, Iván Gómez-Mestre, Christer Brönmark, Thomas Lakowitz, Johan Hollander, Wilco C.E.P. Verberk, David T. Bilton, Anssi Laurila, Beatrice Lindgren, Ane T. Laugen, Marjan De Block, Mark A. McPeek and Robby Stoks for their direct contributions to the data base.

